# Synaptic plasticity of prefrontal long-range inhibition regulates cognitive flexibility

**DOI:** 10.1101/2025.06.27.662040

**Authors:** Xiyu Zhu, Lara L. Hagopian, Kira E. Wallquist, Vikaas S. Sohal

## Abstract

While glutamatergic synaptic plasticity is believed to be a fundamental mechanism mediating learning, the behavioral significance of plasticity at cortical GABAergic synapses remains less well understood. Furthermore, despite recent discoveries of long-range projections from neocortical GABAergic neurons, details about how they function are also sparse. Here we combine behavioral optogenetics with patch-clamp electrophysiology to link plasticity at long-range GABAergic synapses with higher-order cognitive functions. Specifically, learning extradimensional rule shifts potentiates callosal GABAergic synapses from prefrontal parvalbumin-expressing (PV) neurons onto corticothalamic neurons. Disrupting this potentiation by inhibiting callosal PV terminals during rule shifts induces perseveration, whereas reinstating this potentiation with subsequent gamma-frequency callosal PV terminal stimulation restores flexible behavior. This shows how a novel plasticity locus can regulate brain circuits underlying normal cognition and pathological states.

## INTRODUCTION

Cognitive flexibility – the ability to adapt thoughts and behaviors in response to changes in the environment – depends critically on the prefrontal cortex (PFC) (*1*). Impairments in this function are a hallmark of schizophrenia, contributing significantly to long-term disability and often remaining refractory to existing treatments (*2, 3*). PFC parvalbumin-expressing (PV) neurons are a key source of circuit inhibition believed to generate network dynamics that promote flexible cognition (*4, 5*). However, our understanding of the precise circuit mechanisms through which PV+ inhibitory neurons support cognitive flexibility remains limited. Elucidating these mechanisms is necessary both to understand how PFC implements computations underlying cognitive and behavioral flexibility, and to develop circuit-based therapies that restore flexibility in schizophrenia and other disorders.

Synaptic plasticity is a well-established mechanism for learning (*6–9*), but compared to glutamatergic synapses, experience-dependent synaptic plasticity at GABAergic synapses, including those formed by cortical PV neurons, is less well understood (*10*). Slice electrophysiology studies have successfully induced bidirectional changes in synapses from PV neurons (*11–14*), indicating an inherent capacity for plasticity. Many studies have shown that salient experiences can induce cortical inhibitory circuit plasticity by altering the intrinsic properties of inhibitory interneurons and/or their excitatory inputs (*15–23*). Learning stereotyped movements or sensory associations has been shown to alter synapses formed by somatostatin interneurons in motor and barrel cortex, respectively (*8, 24*), and auditory learning is associated with broad changes in spontaneous IPSCs in auditory cortex (*25*). Chronic manipulations that affect excitation-inhibition balance can lead to homeostatic changes in cortical inhibition, including synapses from PV neurons (*26–28*). Modeling work also suggests that inhibitory synaptic plasticity could explain observed patterns of activity during hippocampal replay (*29*). However, there is still a paucity of direct evidence for experience-driven plasticity at GABAergic synapses formed by neocortical PV neurons, and it is unclear how such plasticity might contribute to cognitive flexibility or other higher-order cognitive functions mediated by prefrontal cortex.

In this context, our laboratory has previously shown that within the mouse PFC, PV neurons play a critical role in cognitive flexibility by synchronizing across the hemispheres at gamma-frequencies (∼40 Hz) (*30*). Notably, whereas inhibitory neurons in the PFC and other regions of neocortex have long been assumed to be locally-projecting interneurons, we found that this interhemispheric gamma synchrony depends on a novel class of long-range connections that originate from PV neurons and target pyramidal neurons in the contralateral PFC, particularly those which project to mediodorsal (MD) thalamus (*31*). Remarkably, these callosal PV (cPV) projections appear to bidirectionally regulate cognitive flexibility: optogenetic inhibition of cPV projections during a cognitive flexibility task results in perseverative behaviors and impaired gamma synchrony that persist on subsequent days. Conversely, subsequent gamma-frequency (40Hz) stimulation of these projections elicits an enduring rescue of perseverative behavior (*31*). These effects parallel our earlier finding that synchronous 40 Hz stimulation of bilateral PFC PV neurons also restored cognitive flexibility in mutant mice which model aspects of schizophrenia (*30, 32*). The long-lasting changes in cognitive flexibility we observed after targeting cPV terminals raise the possibility that these manipulations may act by modulating endogenous plasticity mechanisms at this synaptic locus.

## METHODS

### Mice

All animal procedures were conducted in accordance with NIH guidelines and approved by the Administrative Panels on Laboratory Animal Care at the University of California, San Francisco. Both male and female C57BL/6J mice (Jackson Laboratory) were used in this study. Male and female mice were group-housed (2–5 per cage) with standard bedding in a temperature-controlled environment (22–24°C), under a 12 h light/12 h dark cycle, with ad libitum access to food and water until the onset of rule-shift experiments. Experiments were conducted using *PV-Cre* and *PV-Cre:Ai14* transgenic lines. All behavioral and slice electrophysiology experiments were conducted when mice were 11–18 weeks old. All behavioral testing and recordings were performed using age-matched littermates with no more than a 7-day age difference between animals.

### Surgery

All surgeries were performed on mice aged 4–6 weeks. Mice were anesthetized with isoflurane (3% induction, 1–2% maintenance in 95% oxygen) and positioned in a stereotaxic frame (David Kopf Instruments), with body temperature maintained using a heating pad. After exposing the skull, stereotaxic alignment was confirmed using bregma and lambda. Viral injections were delivered at 100 nL/min using a 10 μL Nanofil syringe (World Precision Instruments) fitted with a 35-gauge beveled needle and controlled by a syringe pump (UMP3; World Precision Instruments). The needle remained in place for ≥5 min post-infusion to ensure diffusion before being slowly withdrawn. Following surgery, mice recovered on a heated pad until ambulatory and were then returned to their home cages. Unilateral injection sides were counterbalanced across animals.

The mPFC coordinates for injections (in mm, relative to Bregma) were: +1.7 anterior-posterior (AP), + or – 0.3 mediolateral (ML), and –2.25 dorsoventral (DV). Coordinates for medial dorsal thalamus (MD) were: +1.7 AP, + or – 0.35 ML (contralateral to PFC injections), and –3.5 DV.

For the experiment involving different rule learning paradigms (Fig. 1), *PV-Cre* mice received an injection of AAV5-EF1α-DIO-ChR2-eYFP (600 nL; UNC Vector Core) into the PFC, and AAVrg-CAG-tdTomato (200 nL; Addgene) into the contralateral MD. For experiments involving optogenetic manipulation of callosal PV (cPV) axon terminals (Fig. 2-3), a mixture of AAV5-EF1α-DIO-ChR2-eYFP (600 nL) and AAV5-EF1α-DIO-eNpHR3.0-mCherry (600 nL; UNC Vector Core) was injected into one hemisphere of the mPFC in *PV-Cre* mice, and AAVrg-CAG-tdTomato (200 nL; Addgene) was injected into the contralateral MD. A mono fiber-optic implant (core/outer diameter: 200/240 μm, NA 0.22; Doric Lenses, MFC_200/240-0.22_2.0mm_FLT) was inserted into the contralateral mPFC at coordinates: +1.7 AP, + or –0.35 ML, –1.8 DV. Implants were affixed to the skull using Metabond Quick Adhesive Cement (Parkell). For electrophysiological recordings of intrinsic PV neuron properties after cPV (Fig. S4), *PV-Cre:Ai14* mice received an injection of AAV5-EF1α-DIO-eNpHR3.0-mCherry (500 nL; UNC Vector Core) into one PFC hemisphere. In the contralateral PFC, AAVrg-DIO-eYFP (500 nL; Addgene) was injected, and a mono fiber-optic implant (200/240 μm diameter; Doric Lenses) was placed at: AP –1.7 AP, + or –0.35 ML, and –1.8 DV. All behavioral and electrophysiological recordings were performed at least 7 weeks after virus injection to allow for sufficient expression of viral constructs in axon terminals. Viral titers (in vg/mL) were as follows: AAV5-EF1α-DIO-ChR2-eYFP: 2.3–4.5 × 10¹²; AAV5-EF1α-DIO-eNpHR3.0-mCherry: 2.3–5.3 × 10¹²; AAVrg-CAG-tdTomato: 2.1 × 10¹³; AAVrg-DIO-eYFP: 2.4 × 10¹³.

**Fig. 1.**
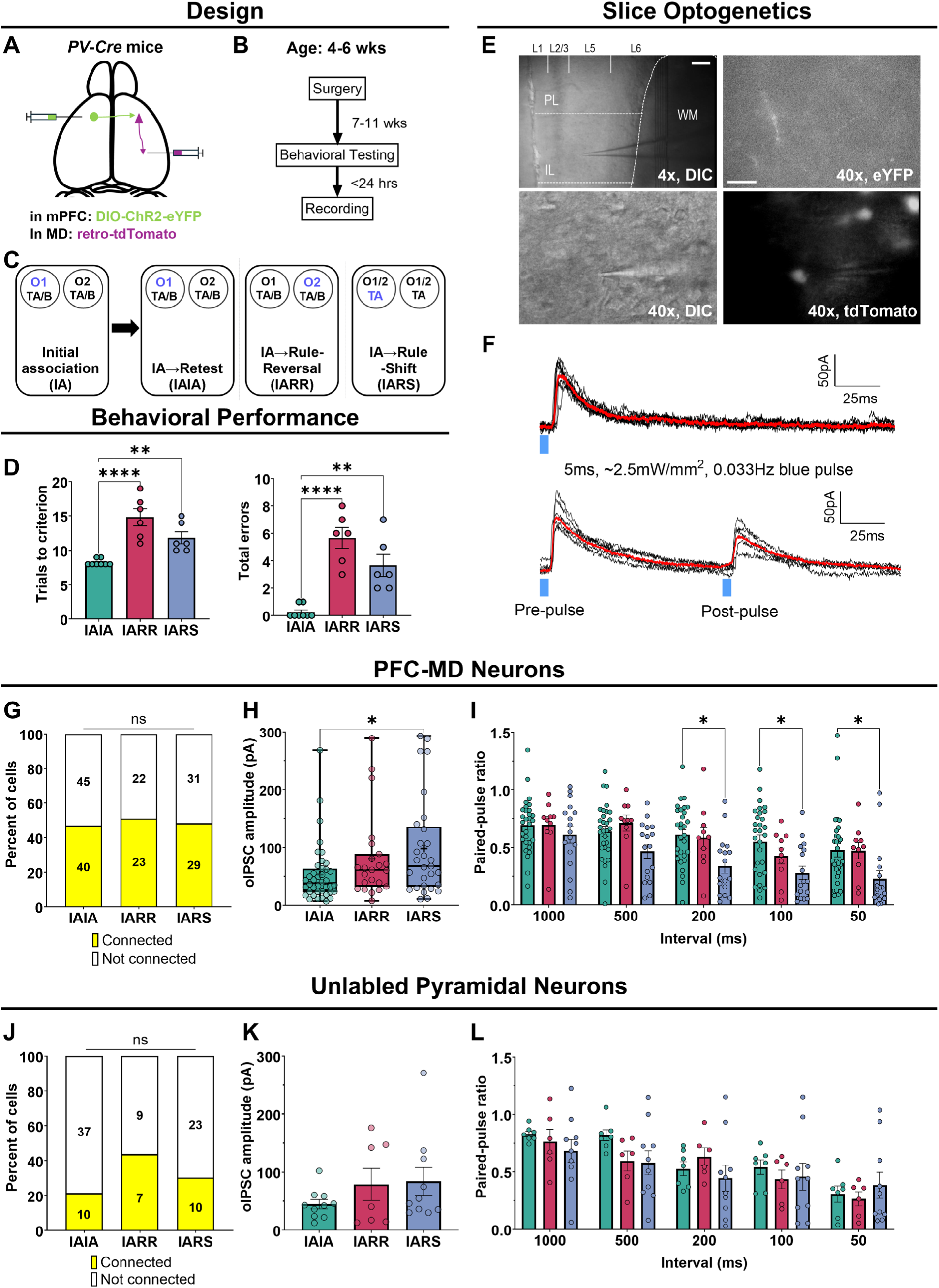
Rule-shift learning strengthens callosal PV synapses onto PFC-MD neurons. **(A)** Experimental design (surgery): in *PV-Cre* mice, PFC PV neurons in one hemisphere were labeled using AAV5-DIO-ChR2-eYFP, and PFC-MD neurons were retrogradely labeled by injecting AAVrg-CAG-tdTomato (in the contralateral hemisphere). **(B)** Experimental timeline: 7-11 weeks after surgery, mice were tested in one of three possible learning paradigms, followed by ex vivo whole-cell recordings within 24 hours of the completion of behavioral testing. **(C)** Task structure: briefly, each trial mice choose to dig in one of two bowls to find a food reward. Bowls are each scented with a different odor (O1 or O2) and filled with a different textured digging medium (TA or TB). Mice must first learn an initial association (IA) between food reward and one sensory cue (e.g., O1). Once a learning criterion (8 correct out of 10 consecutive trials) is reached, the rule either repeats (IAIA) or undergoes an intradimensional reversal (IARR) or extradimensional rule-shift (IARS). The cue associated with reward is indicated in purple. **(D)** IARS (n = 6) and IARR (n = 6) mice required more trials (<u>left</u>) to reach the second learning criterion and made more errors <u>(right)</u> (n = 8; one-way ANOVA). **(E)** Representative images showing recordings in PFC contralateral to AAV injection. <u>Top left</u>: Low-magnification image (scale bar = 50 μm) of a deep layer recording site near the prelimbic/infralimbic border. <u>Top right</u>: High-magnification image of ChR2-eYFP in cPV terminals in L5 (scale bar = 12.5 μm). <u>Bottom</u>: High-magnification DIC images showing a patched neuron which expresses tdTomato. **(F)** Example voltage clamp recordings (at 0 mV) of postsynaptic responses to optogenetic stimulation of cPV terminals (black traces: single responses; red traces: averaged responses). **(G)** cPV→PFC-MD connection probability did not differ between learning paradigms (χ^2^ test). White and yellow bars indicate the proportion of unconnected vs. connected cells, respectively; overlaid numbers indicate actual cell numbers. **(H)** oIPSC amplitudes in connected PFC-MD neurons from IARS mice (98 ± 16 pA; N = 29 cells from 6 mice) were increased compared to IAIA mice (53 ± 8 pA; N = 40 cells from 8 mice); oIPSC amplitudes from IARR mice (80 ± 15; N = 23 cells from 6 mice) were not significantly different from either group (Kruskal-Wallis test, Dunn’s post hoc test, p<0.05). **(I)** The paired pulse ratio (PPR) was lower in PFC-MD neurons from IARS (N = 17 cells from 6 mice) vs. IAIA (N = 30 cells from 8 mice) groups at paired-pulse interval: 200, 100, and 50 ms (repeated measures two-way ANOVA, Bonferroni’s post hoc test; main effect, learning: p<0.01). No changes were observed in the IARR group (n = 10 cells from 6 mice). **(J-L)** Similar to G-I but for unlabeled pyramidal neurons. Error bars indicate +/-SEM. Box-and-whisker plots are used for nonparametric distributions, and show medians (center line), means (‘+’), interquartile ranges (box border), whisker (min/max) indicated. * p < 0.05, ** p < 0.01, *** p<0.001, **** p<0.0001. See figure S1 and table S1 for additional results and detailed statistics.

**Fig. 2.**
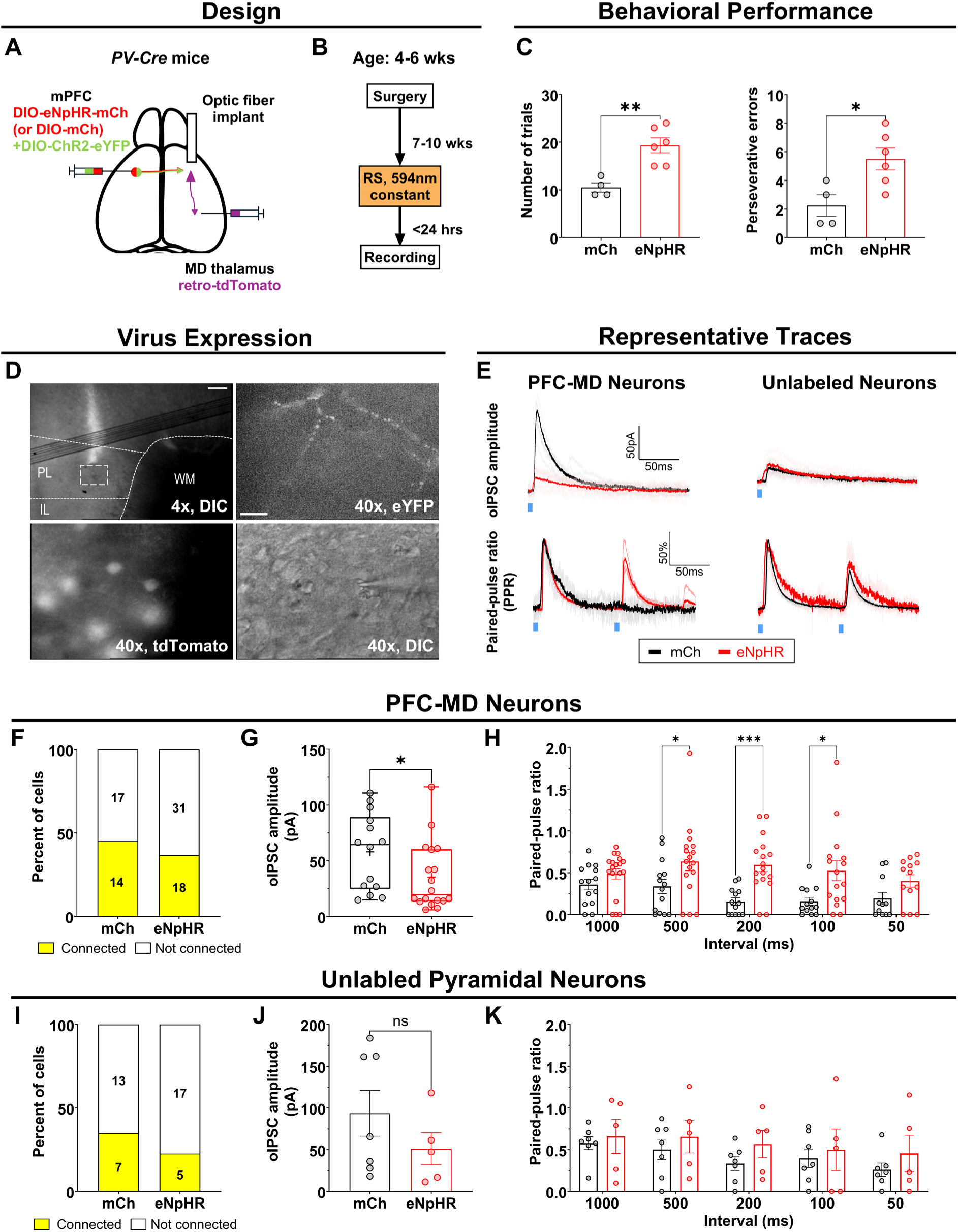
Inhibiting cPV terminals during rule-shift learning causes perseveration and disrupts cPV→PFC-MD synapses. **(A)** Experimental design (surgery): we injected *PV-Cre* mice in one mPFC with AAV5-DIO-ChR2-eYFP mixed either with AAV5-DIO-eNpHR-mCherry (eNpHR) or AAV5-DIO-mCherry (mCh). A 0.2mm diameter optic fiber was implanted in the contralateral mPFC, targeting deep layers. AAVrg-CAG-tdTomato was injected in the MD thalamus, ipsilateral to the fiber implant. **(B)** Experimental timeline: 7-10 weeks after surgery, mice learned an IA followed by an RS. During the RS portion, light was delivered for optogentic inhibition (continuous, 594nm, 5mW). We prepared slices and performed voltage clamp recordings <24 hrs after completing behavior. **(C)** cPV terminal inhibition increased trials to criterion (left) and perseverative errors (right) in eNpHR+ mice (n = 6) compared to controls which expressed mCh only (n = 8) (unpaired t-tests). **(D)** Representative images of whole-cell recordings in contralateral mPFC. <u>Top left</u>: low-magnification image (scale bar = 50 μm) of an implant at the PL/IL border. <u>Top right</u>: high-magnification image (scale bar = 12.5 μm) of ChR2-eYFP in cPV terminals. <u>Bottom</u>: high-magnification image of a patch pipette targeting a tdTomato+ PFC-MD neuron. **(E)** Representative voltage clamp traces of oIPSCs recorded from PFC-MD neurons in eNpHR mice (light red/red) compared to mCh control (gray/black). **(F)** cPV inhibition did not alter the cPV→PFC-MD connection probability (p = 0.45, χ^2^ test). **(G)** oIPSC amplitudes were increased in PFC-MD neurons from eNpHR+ mice (58 ± 9 pA, N = 14 cells from 4 mice) vs. control expressing mCh only (35 ± 7 pA, N = 18 cells from n = 6 mice; Mann-Whitney test). **(H)** cPV→PFC-MD synapses in eNpHR mice showed increased PPR for ISIs of 500, 200 and 100ms (repeated measure two-way ANOVA, Bonferroni’s post hoc; main effect, inhibition: p<0.001). **(I-K)** Similar to F-H for unlabeled pyramidal neurons. See figure S2 and table S2 for additional results and detailed statistics.

**Fig. 3.**
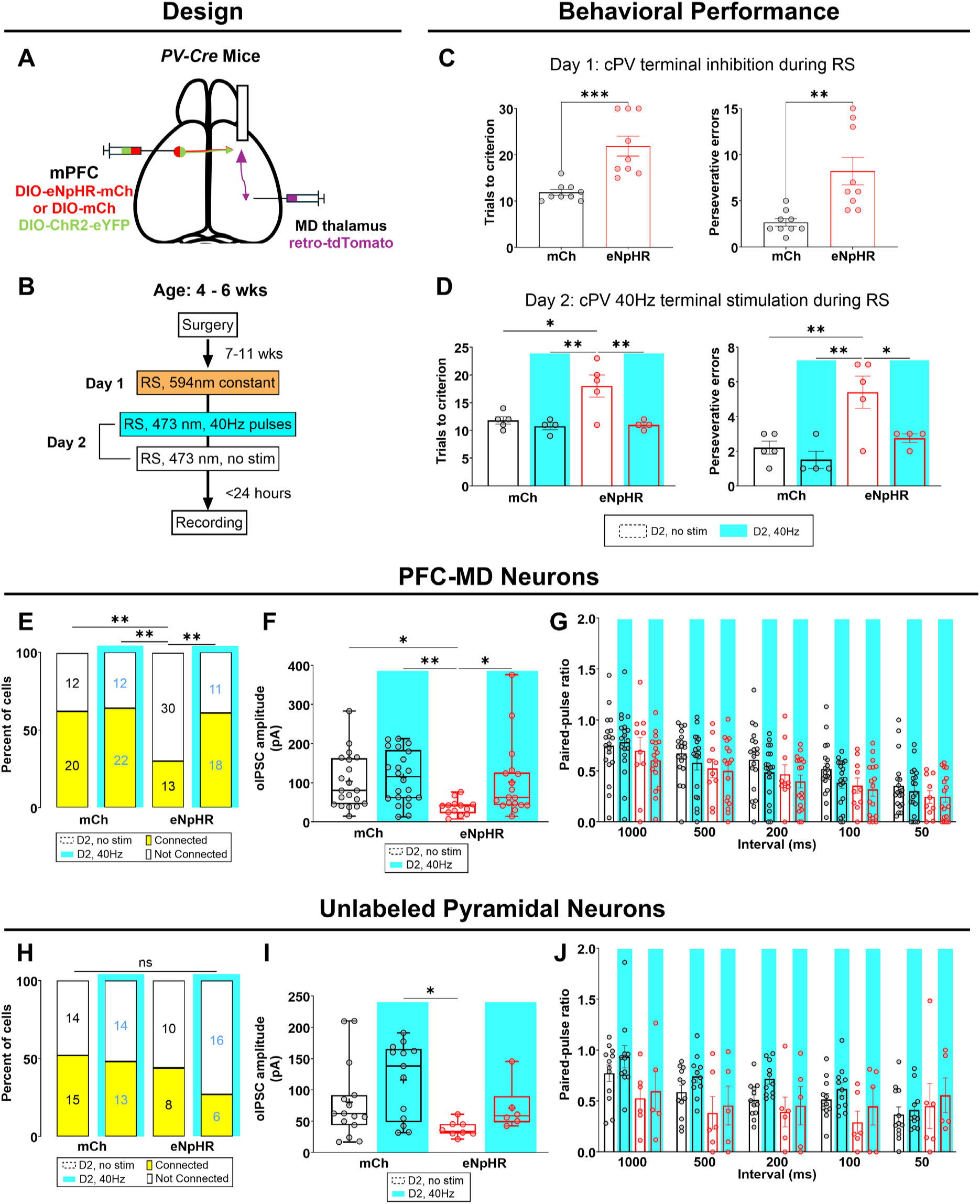
cPV terminal inhibition-induced perseveration and and cPV→PFC-MD synaptic deficits are reversed by subsequent 40Hz stimulation. **(A)** Experimental design (surgery): we injected *PV-Cre* mice in one mPFC with AAV5-DIO-ChR2-eYFP mixed either with AAV5-DIO-eNpHR-mCherry (eNpHR) or AAV5-DIO-mCherry (mCh). A 0.2mm diameter optic fiber was implanted in the contralateral mPFC, targeting deep layers. AAVrg-CAG-tdTomato was injected in the MD thalamus, ipsilateral to the fiber implant. **(B)** Experimental timeline: 7–11 weeks after surgery, mice performed rule-shift learning on two consecutive days. Day 1: cPV projections were inhibited during the rule-shift portion of the task (continuous 594nm light, 5 mW). Day 2: cPV projections were either stimulated at 40 Hz (5ms flashes of 473nm laser, 2.5 mW) or unstimulated. Acute PFC slices were collected within 24 hours for whole-cell recordings. **(C)** Behavioral effects of cPV inhibition on Day 1: cPV terminal inhibition increased trials to criterion (left) and perseverative errors (right) in eNpHR+ mice (n = 9), compared to mCh-only controls (n = 9) (unpaired *t* test). **(D)** On Day 2 mice in each group either receive 40 Hz cPV stimulation (D2, 40 Hz’; blue) or are unstimulated (‘D2, no stim’; no shading). Rule shift learning deficits induced by eNpHR-mediated optogenetic inhibition on Day 1 persist on Day 2 in the ‘no stim’ group but are rescued in the ‘40 Hz’ group (two-way ANOVA with Bonferroni post hoc; trials to criterion – effect of eNpHR: p < 0.05, effect of 40 Hz stimulation: p < 0.01, interaction: p < 0.05; perseverative errors – effect of eNpHR, p < 0.01; 40 Hz stimulation, p < 0.05; interaction, p < 0.05). **(E)** The cPV→PFC-MD connection probability was significantly reduced in eNpHR, ‘D2, no stim’ mice compared to mCh controls, but rescued by 40 Hz stimulation (χ^2^ test). **(F)** cPV inhibition led to reduced oIPSC amplitude in PFC-MD neurons from eNpHR+/no stim mice (37 ± 5 pA; N = 13 neurons from 5 mice) compared to mCh-only/no stim controls (103 ± 16 pA; N = 20 neurons from 5 mice). oIPSC amplitude was rescued in eNpHR+/40 Hz mice (102 ± 22 pA; N = 18 neurons from 4 mice; Kruskal–Wallis test with Dunn’s post hoc). **(G)** Neither cPV inhibition nor 40 Hz stimulation affected the PPR of cPV→PFC-MD synapses. **(H-J)** Similar to E-G for unlabeled pyramidal neurons. oIPSC amplitudes were significantly reduced in neurons from eNpHR/no stim mice (38 ± 4 pA; N = 8 neurons from 5 mice) compared to the mCh/40 Hz group (116 ± 17 pA; N = 13 neurons from 4 mice). See figure S3 and table S3 for additional results and detailed statistics.

### Rule-shift and rule-reversal learning tasks to assess cognitive flexibility

Rule-shift and rule-reversal tasks assessing cognitive flexibility were conducted as previously described (*30, 31*), with minor modifications. Briefly, these tasks were performed in the home cages, requiring mice to attend to one relevant sensory cue (an odor or texture within the digging media) to find a hidden food reward (20 mg food pellets; Bio-Serv). Each trial, mice were presented with two bowls, each containing a compound cue composed of one odor and one digging medium. Only one bowl was baited with the reward. A choice was defined by sustained digging (>0.5 s) in one of the bowls, after which the non-chosen bowl was immediately removed. Stimuli were presented using two identical small animal food bowls (All Living Things Nibble Bowls, PetSmart). Olfactory cues were created using ground dried spices (garlic or coriander powder; McCormick), and sensory/visual cues were provided by unscented digging media (Mosser Lee White Sand Soil Cover or Natural Integrity Clumping Clay Cat Litter). To standardize texture and odor intensity, digging media were mixed with the spices and food pellet powder (to mask the smell of the reward) at a concentration of 0.1% by volume.

One week prior to testing, mice were singly housed and habituated to a reverse light/dark cycle. Food intake was restricted to maintain body weight at 80–85% of ad libitum levels. Mice then underwent a single habituation session consisting of ten consecutive trials. In each trial, one bowl was randomly baited. By the end of this habituation session, all mice would readily dig in bowls to retrieve the food reward. No additional training was conducted prior to testing.

The first phase of the task was the *initial association* (IA), during which mice were presented with two bowls containing compound cues. Only one cue reliably predicted the location of the hidden reward. IA continued until mice achieved 8 correct choices in 10 consecutive trials. Upon reaching this criterion, mice proceeded to the next task phase depending on the specific learning paradigm. In a *rule-shift* (RS), the predictive cue was switched to the other sensory modality (i.e., from odor to texture or vice-versa). In a *rule-reversal* (RR), the reward-predictive cue was reversed to the other stimulus from the same modality (e.g., from garlic to coriander). We also tested some mice using a paradigm in which the IA rule remained unchanged. In this case, mice were required to complete an additional 8 out of 10 correct trials to complete the second IA phase. Only one task session (e.g., learning of an IA and an RS) was performed per day. Initial IA cues were randomly assigned and counterbalanced across animals for cue dimensions (i.e., some animals started with an odor-based rule, others with a texture-based one). For multi-day tasks (Fig. 3), the IA cue on the second day was always the cue that had been shifted to in the previous session, to control for potential long-lasting associations (i.e., the IA cue on Day 2 was the same as the RS cue from Day 1). In the rule-shift task (Fig 2 and 3), errors in which mice continued responding according to the IA cue were classified as *perseverative errors*. All other incorrect responses were considered *random errors*. During behavioral testing, experimenters were blind to group identity whenever possible. Note that, blinding was not possible for the mice which did or did not receive 40Hz optogenetic stimulation (Day 2 of the experiment in Fig. 3).

### *In vivo* optogenetic inhibition of cPV projections during rule-shift task

For optogenetic inhibition of cPV terminals using eNpHR (Fig. 2–3), a 594 nm orange laser (OEM Laser Systems) was coupled to the implanted mono fiber-optic cannula via a zirconia sleeve and a 200-μm-diameter mono fiber-optic patch cord (all components from Doric Lenses). Laser power at the implant tip was adjusted to 5mW. For optogenetic activation of cPV terminals using ChR2, a 473 nm blue laser (OEM Laser Systems) was similarly coupled to the mono fiber-optic implant. Light power at the tip was adjusted to 2.5mW. A function generator (Agilent 33500B Series Waveform Generator) was used to drive the laser, delivering 40 Hz trains of 5ms pulses. In both protocols, laser stimulation was applied exclusively during the rule-shift phase of the task, as previously described (*31*).

### Slice electrophysiology

#### Slice preparation and recording conditions

Within 24 hours of behavioral testing, mice were anesthetized with isoflurane and rapidly decapitated for brain extraction. Coronal brain slices (250μm thick) containing the mPFC were prepared using a vibratome (VT1200S, Leica Microsystems) in a chilled sucrose-based slicing solution in which Na⁺ was replaced by sucrose. Slices were then incubated in warmed artificial cerebrospinal fluid (ACSF) at 30–33 °C for 30 minutes, followed by at least 30 minutes hour at room temperature before electrophysiological recordings. The external ACSF used for holding slices contained (in mM): 125 NaCl, 25 NaHCO₃, 3 KCl, 1.25 NaH₂PO₄, 2 MgCl₂, 2 CaCl₂, 10 D-glucose, 1.3 ascorbic acid, and 3 sodium pyruvate. The recording ACSF contained: 125 NaCl, 25 NaHCO₃, 3 KCl, 1.25 NaH₂PO₄, 1 MgCl₂, 2 CaCl₂, 10 D-glucose.

#### Optogenetically-evoked IPSC (oIPSC) recording

Whole-cell voltage-clamp recordings were performed from layer 5/6 pyramidal neurons in the mPFC (*33, 34*). In experiments with implanted optical fibers (Fig. 2-3), recordings were made below the fiber tip (corresponding to AP 1.98–1.34 mm, Paxinos and Franklin Mouse Brain Atlas, 2nd ed). Only slices with implants accurately targeting the prelimbic (PL) cortex were included. Recording sites were restricted to slices containing the most ventral visible implant mark and its immediate neighbors. Pyramidal neurons were identified based on morphology (e.g., prominent apical dendrite) under IR-DIC optics (Olympus BX51WI). PFC–MD-projecting neurons were identified by retrograde tdTomato labeling from virus injections into the MD. Recordings were performed using a Multiclamp 700A amplifier (Molecular Devices) and patch pipettes (2–4 MΩ) pulled from borosilicate glass (Sutter Instruments). The internal solution contained (in mM): 130 Cs-methanesulfonate, 10 CsCl, 10 HEPES, 4 NaCl, 7 phosphocreatine, 0.3 Na-GTP, 4 Mg-ATP, and 2 QX-314 Br (pH 7.3, adjusted with CsOH). oIPSC recordings began at least 5 minutes after break-in. Cells were held at 0 mV to isolate inhibitory currents. Series resistance (10–20 MΩ) and pipette capacitance were compensated; recordings were discarded if resistance exceeded 25 MΩ. Optogenetic stimulation was generated by a 480 nm LED (Lambda FLED; Sutter Instrument) and delivered through a 40x objective. ChR2-expressing callosal PV (cPV) axons were stimulated with 480 nm light (5-ms pulses at 2.5 mW). For each cell, 10 sweeps (2 s duration, 30 s inter-sweep interval) were used to assess oIPSC amplitude and connectivity. Cells were defined as connected if they showed a failure rate < 50% and mean oIPSC amplitude >5 pA. Connected cells were further tested for paired-pulse ratio (PPR) using five sweeps per interstimulus interval (1000, 500, 200, 100, and 50 ms). Cells with >25% variability in pre-pulse amplitude were excluded. Analysis of oIPSC amplitude and PPR was performed in Clampfit 10.7 by averaging over 3–10 sweeps with time-locked responses. In separate slices, we validated the GABAergic nature of the observed oIPSC events (Fig. S1) using 6-cyano-7-nitroquinoxaline-2,3-dione disodium salt hydrate (CNQX), 2-(3-carboxypropyl)-3-amino-6-(4 methoxyphenyl)-pyridazinium bromide (gabazine), and d-2-amino-5-phosphonopentanoic acid (D-AP5) (all from Tocris). Drugs were prepared as concentrated stock solutions and were diluted in ACSF on the day of the experiment. For all experiments, spontaneous inhibitory postsynaptic currents (sIPSCs) and excitatory postsynaptic currents (sEPSCs) were recorded from a subset of connected PFC-MD cells (Fig. S1-S3). Specifically, 30–90 second gap-free recordings were obtained for analysis. sEPSCs were recorded at a holding potential of –70 mV, starting approximately 5 minutes after establishing the whole-cell configuration and prior to oIPSC recordings. sIPSCs were recorded at 0 mV, approximately 2 min following oIPSC recordings and before PPR measurements. Spontaneous events (amplitude > 10pA) were detected automatically using the Mini Detection package in SimplyFire 0.6.1 software (*35*) and subsequently verified manually.

#### Current-Clamp Recordings of PV Neurons

To assess intrinsic excitability (Fig. S4), current-clamp recordings were performed from tdTomato-labeled PV neurons using an internal solution containing (in mM): 130 K-gluconate, 10 KCl, 10 HEPES, 10 EGTA, 2 MgCl₂, 2 Mg-ATP, and 0.3 Na-GTP (pH 7.3, adjusted with KOH). Responses were elicited by hyperpolarizing and depolarizing current steps (25 or 50 pA increments; −200 to 450 pA; 250 or 800 ms duration). Spiking properties were assessed using a step 100 pA above rheobase. Analysis (Fig. S4A–D) included only regular– and fast-spiking PV neurons; those with high interspike interval variability or low firing rates at 450 pA were excluded (9 of 30 cPV and 5 of 24 local PV cells). No irregular-spiking cPV neurons were observed in the dataset shown in figure S4E–G.

### Data analysis

Statistical analyses were performed using GraphPad Prism. Full details of statistical analyses— including test types, test statistics, sample sizes (N and n), treatment of repeated measures, and corrections for multiple comparisons—are provided in tables S1–S4. All tests were two-sided, and statistical significance was defined as *p* < 0.05. Data are reported as mean ± SEM unless otherwise noted. Normality was assessed using the Shapiro–Wilk test when appropriate. For two-group comparisons, unpaired two-tailed *t*-test (or Mann Whitney U test) was used. For comparisons involving more than two groups, one-way or two-way ANOVA (or their nonparametric equivalents) was applied, followed by appropriate post hoc corrections (e.g., Bonferroni) where applicable. Repeated-measures ANOVA was used for within-subject comparisons.

## RESULTS

### Learning an extradimensional rule shift potentiates cPV-mediated inhibition onto contralateral PFC-MD neurons

As in prior studies from our laboratory and others (*30–32, 36–39*), we assessed cognitive flexibility using a digging-based ‘rule-shifting’ task, adapted from the classical attention set-shifting paradigm in rats (*40*) (see Materials and Methods). In this task, mice chose between two bowls on each trial to find a hidden food reward. Each bowl contains different odor (garlic or coriander) and texture cues (sand or litter) within the digging media, and after mice learn an ‘initial association’ (IA) based on one cue, the cue-reward association undergoes an extradimensional ‘rule shift’ (RS) from an odor based cue (e.g., rewarded cue = garlic) to one based on a texture (e.g., rewarded cue = sand) or vice-versa. We hypothesized that this rule shift may be a salient experience sufficient to potentiate cPV synapses. To test this, we selectively labeled cPV projections by unilaterally injecting AAV-EF1a-DIO-hChR2(H134R)-eYFP into the medial prefrontal cortex (mPFC) of *PV-Cre* mice. Simultaneously, we retrogradely labeled PFC neurons which project to MD thalamus (PFC-MD neurons) by injecting AAVrg-CAG-tdTomato into the contralateral MD (Fig. 1A). After 7-11 weeks of virus expression, mice were assigned to one of three learning paradigms (Fig. 1B), all beginning with learning of an IA. After mice successfully learned the IA, we either retested mice on the IA rule (‘IAIA’; e.g., garlic→garlic), performed an intradimensional rule reversal (‘IARR’; e.g., garlic→coriander), or performed an extradimensional rule shift (‘IARS’; e.g., garlic→sand) (Fig. 1C). Whole-cell patch-clamp recordings were conducted <24 hours after the completion of behavior to study the effects of these different paradigms on cPV synapses (Fig. 1B).

As expected, both IARS and IARR mice required slightly more trials (and made slightly more errors) than IAIA controls before reaching the learning criterion for the second rule (Fig. 1D). There were no differences between these groups during IA learning (Fig. S1A-B). In acute mPFC slices prepared <24 hours after completing behavioral testing, we observed ChR2-eYFP-labeled cPV terminals in layers 5 and 6 of the mPFC contralateral to the site of virus injection. We made whole-cell patch-clamp recordings from retrogradely labeled tdTomato+ PFC-MD neurons as well as from neighboring unlabeled pyramidal neurons (Fig. 1E). Optogenetic stimulation using 5ms blue LED pulses evoked inhibitory postsynaptic currents (oIPSCs) in ∼50% of PFC-MD neurons and a smaller fraction of unlabeled pyramidal neurons (Fig. 1F). Consistent with our prior findings (*31*), we confirmed that oIPSCs evoked by cPV stimulation were completely blocked by the GABA-A receptor antagonist gabazine, but unaffected by glutamatergic antagonists (Fig. S1C-D). Connectivity, i.e., the fraction of PFC-MD neurons with detectable oIPSC responses to cPV stimulation, did not differ between the IAIA, IARS, and IARR groups (Fig. 1G). However, oIPSC amplitudes were significantly larger in the IARS group compared to IAIA controls, while the IARR group showed intermediate responses that were not significantly different from either IAIA or IARS (Fig. 1H). We also observed a similar trend in unlabeled pyramidal neurons, but this did not reach statistical significance (Fig. 1K). Similar to local PV synapses (*41, 42*), most cPV→PFC-MD synapses exhibited strong paired-pulse depression (PPR < 1). Notably, PPR was significantly reduced in the IARS group, relative to IAIA controls, consistent with a presynaptic locus for synaptic potentiation (Fig. 1I). The frequencies and amplitudes of spontaneous excitatory (sEPSC) and inhibitory post-synaptic currents (sIPSC) in connected PFC-MD neurons did not differ between groups (Fig. S1E-J), suggesting a specific potentiation of cPV→PFC-MD synapses, rather than a nonspecific enhancement of all excitatory or inhibitory synaptic inputs. Together, these findings reveal that rule-shift learning selectively potentiates cPV outputs onto PFC-MD neurons.

### Inhibiting cPV projections during rule-shift learning causes perseveration and weakens cPV**→**PFC-MD synapses

Since rule-shift learning normally strengthens cPV→PFC-MD synapses, we next investigated whether presynaptic cPV activity during rule-shift learning is necessary for this plasticity. For this, we injected *PV-Cre* mice in one mPFC with AAV5-EF1a-DIO-hChR2(H134R)-eYFP mixed with either AAV5-EF1a-DIO-eNpHR3.0-mCherry (eNpHR; experimental group) or AAV5-EF1a-DIO-mCherry (mCh; controls). In the contralateral mPFC, we implanted an optical fiber targeting layer 5 of prelimbic cortex to enable optogenetic inhibition of cPV projections during rule-shift learning. We also injected AAVrg-CAG-tdTomato into the MD to retrogradely label PFC-MD neurons (Fig. 2A). After 7-11 weeks of virus expression, all mice learned an IA followed by a RS; during the RS portion of the task, all mice received light for optogenetic inhibition. This was followed <24hr later by whole-cell recordings in acute slices (Fig. 2B). Consistent with our prior study (*31*), inhibiting cPV projections impaired rule-shifting learning, significantly increasing both the trials needed to reach the learning criterion and perseverative errors in eNpHR+ experimental mice compared to controls (which expressed mCh but not eNpHR) (Fig. 2C). Within 24 hours, we prepared acute PFC slices to record from pyramidal neurons under the implant tip, in the vicinity of ChR2-eYFP-labeled cPV fibers (Fig. 2D), and used 5-ms blue LED pulses to stimulate cPV terminals and elicit oIPSCs (Fig. 2E).

We found that oIPSCs in PFC-MD neurons from eNpHR+ mice (which received cPV terminal inhibition during rule-shift learning) had reduced amplitude and increased PPR compared to PFC-MD neurons from mCh-expressing controls (Fig. 2G-H). A decrease in oIPSC amplitude was also present for unlabeled neurons, but did not reach statistical significance (Fig. 2J). There was no difference between sEPSCs or sIPSCs in connected PFC-MD neurons from eNpHR+ mice vs. those from controls (Fig. S2). Thus, in mice learning an RS, the effects of inhibiting cPV terminals are the exact opposite of the changes normally produced by RS learning (Fig. 1). This indicates that inhibiting presynaptic cPV terminal activity is sufficient to prevent or reverse learning-induced cPV synaptic plasticity, and that the disruption of normal plasticity at this locus tracks the terminal inhibition-induced loss of cognitive flexibility.

### 40 Hz cPV stimulation reverses behavioral and synaptic deficits

We previously found that the persistent perseveration and loss of gamma synchrony caused by inhibiting cPV projections during rule-shift learning can be reversed by subsequent gamma-frequency (40 Hz) cPV terminal stimulation (*31*). To examine how this optogenetic rescue affects cPV synapses, we tested mice on two consecutive days of rule-shift learning, while bidirectionally modulating cPV afferent activity (Fig. 3B). Using the same approach illustrated in Figure 2, we labeled cPV projections with ChR2 plus either eNpHR (experimental group) or mCh only (control group; Fig. 3A). After 7–11 weeks of viral expression, mice underwent the first day of rule-shifting. On this day (‘Day 1’), all mice received light for cPV inhibition during the rule-shift phase. On Day 2, both eNpHR+ and mCh-only (control) mice were further split into two groups: one received 40Hz stimulation of cPV projections (470nm, 5ms, ∼2.5mW; 40Hz stim), while the other performed rule-shifting in the absence of further light delivery (no stim; Fig. 3B). Thus, there were four groups: eNpHR+/40Hz stim, eNpHR+/no stim, mCh+/40Hz stim, and mCh+/no stim. All mice performed 2 days of rule-shifting and expressed ChR2+ in PV neurons unilaterally, in one mPFC.

As expected, cPV terminal inhibition on Day 1 disrupted rule-shift learning, as evidenced by significant increases in the number of trials to criterion and perseverative errors (Fig. 3C). On Day 2, eNpHR+ mice which received 40Hz stim showed a recovery of rule-shift performance, whereas deficits persisted in the eNpHR/no stim group (Fig. 3D). Whole-cell recordings revealed corresponding changes in cPV→PFC-MD synapses (Fig. 3E-G). The strength of cPV synapses was significantly reduced in PFC-MD neurons from the eNpHR/no stim group compared to all three other groups; conversely the oIPSC amplitude in the eNpHR/40Hz stim group was not significantly different from the amplitude associated with the mCh/no stim or mCh/40Hz groups (Fig. 3F). Notably, whereas our previous experiments which measured oIPSCs after one day of rule-shifting found that changes in cPV→PFC-MD oIPSC amplitudes were associated with corresponding changes in the PPR but not in connectivity (Fig. 2F-H), here we observed the opposite pattern: there was no change in PPR between these groups (Fig. 3G), however connectivity (the fraction of PFC-MD neurons with detectable oIPSC responses to cPV stimulation) was significantly reduced in the eNpHR/no stim group compared to all other groups (Fig. 3E). Paralleling the stimulation-induced rescue of oIPSC amplitude, connectivity in the eNpHR/40Hz group was also not different from the mCh/no stim and mCh/40Hz groups. Interestingly, the overall connectivity was significantly higher for the three groups that performed two days of rule-shifting and had normal RS performance on Day 2 (Fig. 3E) compared to the groups from previous experiments that performed only one day of rule-shifting (Fig. 1G, 2F) (63% vs. 47% connectivity, p = 0.029, χ^2^ test). There was also a trend towards weaker cPV oIPSCs in unlabeled pyramidal neurons from the eNpHR/no stim group compared to the other groups (Fig. 3I), although this only reached statistical significance for the mCh/40Hz stim group. Of note, similar to prior experiments we did not observe any changes in sIPSCs or sEPSCs associated with persistent behavioral deficits or rescue (Fig. S3C-F). Thus, we find that gamma-frequency stimulation of cPV terminals, which rescues cognitive flexibility, also restores the strength of cPV→PFC-MD synapses and reverses other synaptic changes caused by cPV terminal inhibition. We also find some possible differences in whether plasticity after 1 vs. 2 days of rule-shifting manifests as altered presynaptic release probability (reflected by PPR) vs. connectivity, suggesting that the passage of time and/or repeated RS learning may induce a shift from functional to structural plasticity.

### Intrinsic properties of callosally-projecting PV neurons are similar to those of local PV interneurons and unaffected by cPV terminal inhibition

Finally, to characterize the intrinsic properties of callosally-projecting PV neurons, compare them to locally-projecting PV interneurons, and evaluate how they are affected by cPV terminal inhibition, we labeled cPV somata by injecting *PV-Cre:Ai14* mice unilaterally with AAVrg-DIO-eYFP in one mPFC (Fig. S4A). After waiting at least 7 weeks post-injection, we made interleaved whole-cell current-clamp recordings from identified callosally-projecting PV (tdTomato+/eYFP+) neurons and neighboring locally-projecting PV (tdTomao+/eYFP-) neurons in acute mPFC slices (contralateral to the site of virus injection). Identified cPV neurons were distributed across cortical layers, predominantly layers 5 and 6, with none detected in layer 1 (Fig. S4C). Although callosally– and locally-projecting PV neurons exhibited largely similar electrophysiological profiles (Fig. S4D), we observed significantly larger spike afterhyperpolarization (AHP) in callosally-projecting PV neurons (table S4), consistent with a prior study of callosally-projecting PV neurons in auditory cortex (*43*). Furthermore, we did not observe any changes in the intrinsic properties of callosally-projecting PV neurons following cPV terminal inhibition (Fig. S4E-G; table S4).

## DISCUSSION

Although synaptic plasticity is generally assumed to underlie learning, the specific synaptic changes which mediate cognitive flexibility remain unclear. When task contingencies change, prefrontal neurons are thought to generate activity patterns which drive adaptive changes in behavioral strategies (*44–46*). Models for storing and updating activity patterns have largely focused on the Hebbian theory of modifying excitatory synapses within recurrent networks to create self-reinforcing neuronal assemblies (*47*). While there has been increasing interest in possible roles of plasticity in GABAergic circuits, it has been challenging to identify specific forms of learning which involve changes in inhibitory synapses formed by neocortical PV neurons. Here, not only do we find that synaptic plasticity downstream of PV neurons plays a key role in cognitive flexibility, but this plasticity actually localizes to long-range GABAergic connections, whose function has been unclear. Specifically, learning an extradimensional rule-shift potentiates GABAergic synapses from prefrontal PV neurons to corticothalamic neurons in the contralateral hemisphere. Disrupting this potentiation, by inhibiting cPV terminals during the rule-shift, prevents learning and creates an enduring perseverative state. Conversely, restoring this potentiation, by delivering gamma-frequency stimulation to cPV terminals during a rule-shift, is associated with a normalization of rule-shift learning. Together these results establish a strong link between cognitive flexibility and the plasticity of prefrontal cPV synapses, and show how plasticity at this locus could regulate PFC-MD connectivity implicated in both normal cognition and pathological conditions such as schizophrenia (*48*).

### Potential involvement of PFC cPV synaptic plasticity in behavioral flexibility

How might cPV plasticity contribute to flexible rule learning? One possibility is that cPV synapses are selectively strengthened on pyramidal neurons that form ensembles and/or representations associated with the initial rule. Thus, potentiation of these cPV synapses could suppress outdated rule representations to prevent perseverative behaviors. This would align with a recently developed model of inhibitory synaptic plasticity in the hippocampus, in which the strengthening of local inhibitory synapses onto specific neurons suppresses activity related to irrelevant cues during sharp-wave ripples (*29*). Another possibility is that cPV→PFC-MD synapses are strengthened more broadly (not just onto specific neurons) in order to alter PFC network dynamics. Thus, increased cPV inhibition might destabilize activity patterns within PFC-MD loops that serve to maintain previously learned associations (*49–51*), promote the emergence of alternative network states that enable the formation of novel associations, and/or facilitate synaptic plasticity at additional loci (possibly by synchronizing pre and post-synaptic activity to gamma rhythms). Thus, the strengthening of cPV inhibition might help promote flexible reorganization of circuit dynamics without directly encoding task-relevant information. This might be similar, at a conceptual level, to how circuit inhibition controls critical period plasticity (*52*).

The deficit we observed in cPV→PFC-MD synaptic strength after cPV inhibition parallels our previous finding that cPV inhibition also persistently disrupts increases in interhemispheric PV gamma synchrony which normally occur during rule-shifts (*31*). Thus, cPV synaptic plasticity may be necessary for generating increases in interhemispheric PV gamma synchrony (*31*). It is also possible that gamma synchrony normally facilitates cPV synaptic potentiation during learning, potentially by aligning presynaptic spikes in callosally-projecting PV neurons with postsynaptic spikes in PFC-MD neurons, and/or by promoting burst firing at frequencies associated with plasticity at PV neuron synapses (*11–13*). These types of mechanisms would be consistent with previous descriptions of activity-dependent plasticity at local PV synapses, although neuromodulatory signals may also contribute (*53, 54*).

### Clinical implications

Although pathological changes in the PFC, including deficits in PV neurons, gamma oscillations, and connectivity with MD thalamus, have long been linked to schizophrenia (*48, 55*), a key challenge in unraveling the pathophysiology of this condition is identifying associated maladaptive states and understanding why they persist. Our findings that cPV terminal inhibition leads to long-lasting deficits in RS learning, gamma synchrony (*31*), and cPV→PFC-MD synaptic strength suggests a plausible pathological cascade: some initial pathogenic insult weakens cPV synapses, compromising the ability of cPV circuits to generate the gamma synchrony required for synaptic potentiation, thus creating an enduring maladaptive state (which would include aberrant regulation of PFC-MD circuitry). Conversely, we find that artificially inducing gamma-frequency activity in cPV projections is sufficient to potentiate them during RS learning, reversing synaptic and behavioral deficits (Fig. 3). This highlights the potential of cPV synapses as a critical node for intervention and suggests that enhancing their plasticity may be a viable therapeutic strategy. Future efforts to molecularly profile cPV neurons – e.g., identify their unique genetic markers, ion channels, or GPCRs – could support the development of novel pharmacological or genetic therapies aimed at restoring synaptic efficacy and plasticity within this circuit. Additionally, since neuroplasticity has been hypothesized to mediate the benefits of existing therapies, e.g., noninvasive brain stimulation and cognitive-behavioral therapy (*6*), our results offers a new framework for understanding and potentially optimizing these interventions.

## Acknowledgments

We acknowledge technical assistance from Lena Shindy and Teagan Bollock.

## Funding

National Institutes of Health grant R01MH129835 (VS), Brain and Behavior Foundation Young Investigator Award (XZ)

**Fig. S1.**
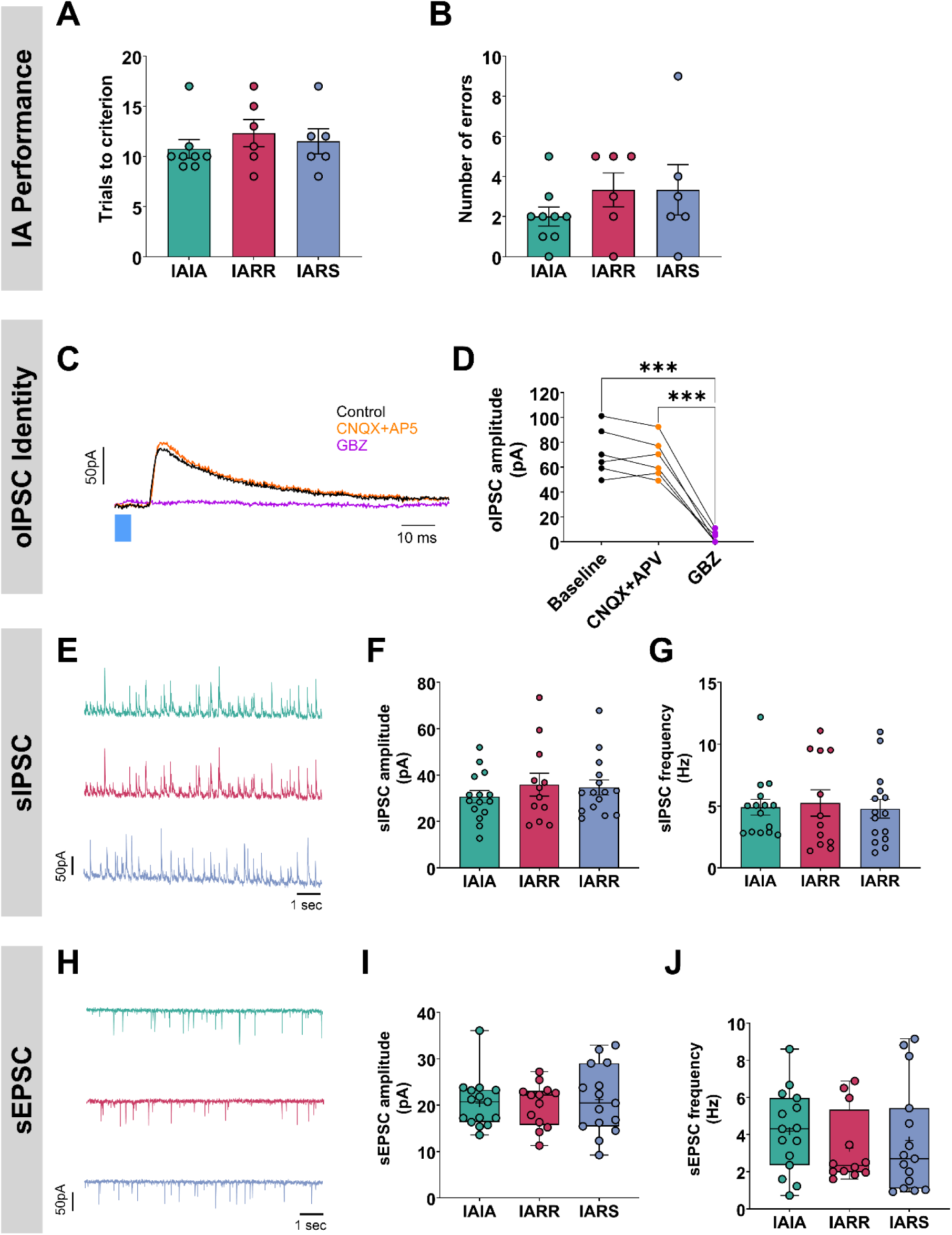
Additional behavioral measures, oIPSC pharmacology, and spontaneous postsynaptic current analyses related to Figure 1. (**A-B**) We did not observe any differences in IA learning during the three different learning paradigms. **(C)** Example recordings from a recipient PFC-MD neuron showing optogenetically evoked IPSCs which persist during application of the glutamatergic antagonists CNQX (10 µM) and APV (50 µM), but are abolished by subsequent application of the GABA-A receptor antagonist gabazine (GBZ, 10 µM). **(D)** Quantification of oIPSC suppression by GBZ but not CNQX and APV (repeated measure ANOVA, p <0.0001; N = 6 cells, 4 PFC-MD neurons, 2 unlabeled pyramidal neurons; n = 2 mice). ***p < 0.001, Bonferroni’s multiple comparisons test. **(E-J)** Spontaneous inhibitory (sIPSC) and excitatory (sEPSC) postsynaptic currents were collected for 30-90 seconds from a subset of connected PFC-MD neurons. **(E**) Example sIPSC recordings. **(F)** Mean sIPSC amplitude from connected PFC-MD neurons were similar among groups (one-way ANOVA, p = 0.56). IAIA: 30 ± 3 pA (N = 15 cells from 8 mice), IARR: 36 ± 5 pA (N = 12 cells from 6 mice), IARS: 35 ± 3 pA (N = 15 cells from 6 mice). **(G)** Mean sIPSC frequencies in connected PFC-MD neurons were similar among groups (p = 0.92) – IAIA: 4.9 ± 0.6 Hz, IARR: 5.2 ± 1.1 Hz, IARS: 4.8 ± 0.8 Hz. **(H)** Example sEPSC recordings. **(I**) Mean sEPSC amplitudes were similar across groups (p = 0.95, Kruskal-Wallis test): IAIA: 20 ± 1 pA, IARR: 21 ± 1 pA, 21 ± 2 pA. **(J)** Mean sEPSC frequencies were similar across groups (p = 0.54) – IAIA: 4.2 ± 0.6 Hz, IARR: 3.3 ± 0.6 Hz, IARS: 3.7 ± 0.8 Hz. Sample size of **(G-J)** is the same as in **(F)**. Bar graph indicates mean ± SEM. Box-whisker plots represent nonparametric dataset determined by Shapiro-Wilk test: line indicates median, box borders indicate IQR, whiskers indicate min/max, and “+” indicates mean.

**Fig. S2.**
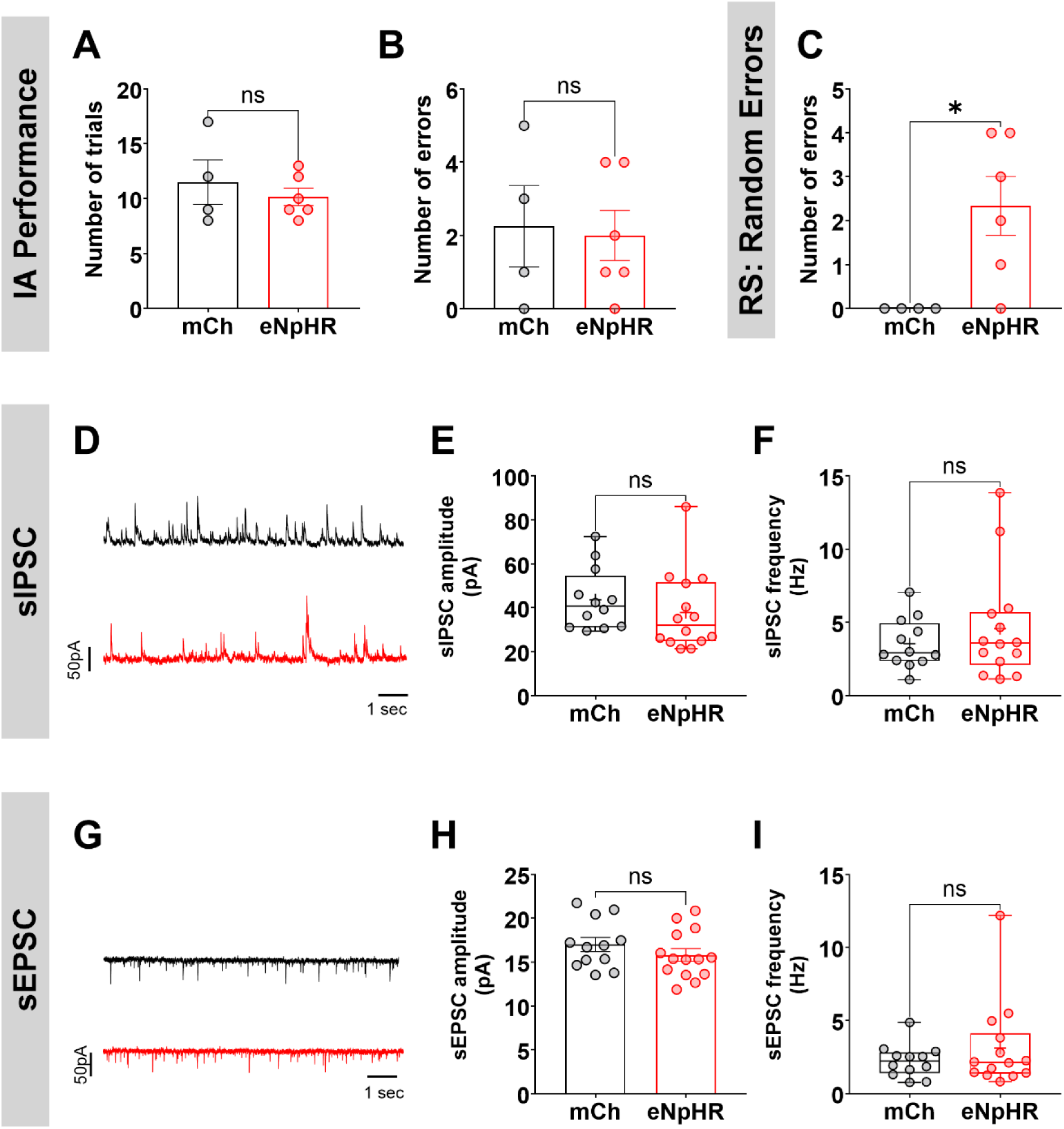
Additional behavioral measures and spontaneous postsynaptic activity recordings related to Figure 2. **(A–B)** Learning of the IA was similar between groups. **(C)** Inhibition of cPV terminals also increased random (i.e., non-perseverative) errors during the rule-shift task, consistent with our previous finding (*31*). **(D–I)** Spontaneous IPSCs and EPSCs were recorded over 30–90 seconds from a subset of connected PFC-MD neurons. Sample sizes: mCh, *N* = 12 cells from 4 mice; eNpHR, *N* = 14 cells from 6 mice. **(D)** Representative sIPSC traces (10 s). cPV inhibition did not significantly alter **(E)** sIPSC amplitude (mCh: 44 ± 4 pA vs. eNpHR: 38 ± 5 pA; Mann Whitney test, p = 0.12) or **(F)** sIPSC frequency (mCh: 3.5 ± 0.5 Hz vs. eNpHR: 4.6 ± 1.0 Hz; Mann Whitney test, p = 0.67). **(G–I)** Similar analyses of spontaneous EPSCs revealed no significant group differences. **(H)** sEPSC amplitude (mCh: 18 ± 1 pA vs. eNpHR: 16 ± 0.7 pA; unpaired *t* test, p = 0.28). **(I)** sEPSC frequency (mCh: 2.2 ± 0.3 Hz vs. eNpHR: 3.1 ± 0.8 Hz; Mann Whitney test, p = 0.78).

**Fig. S3.**
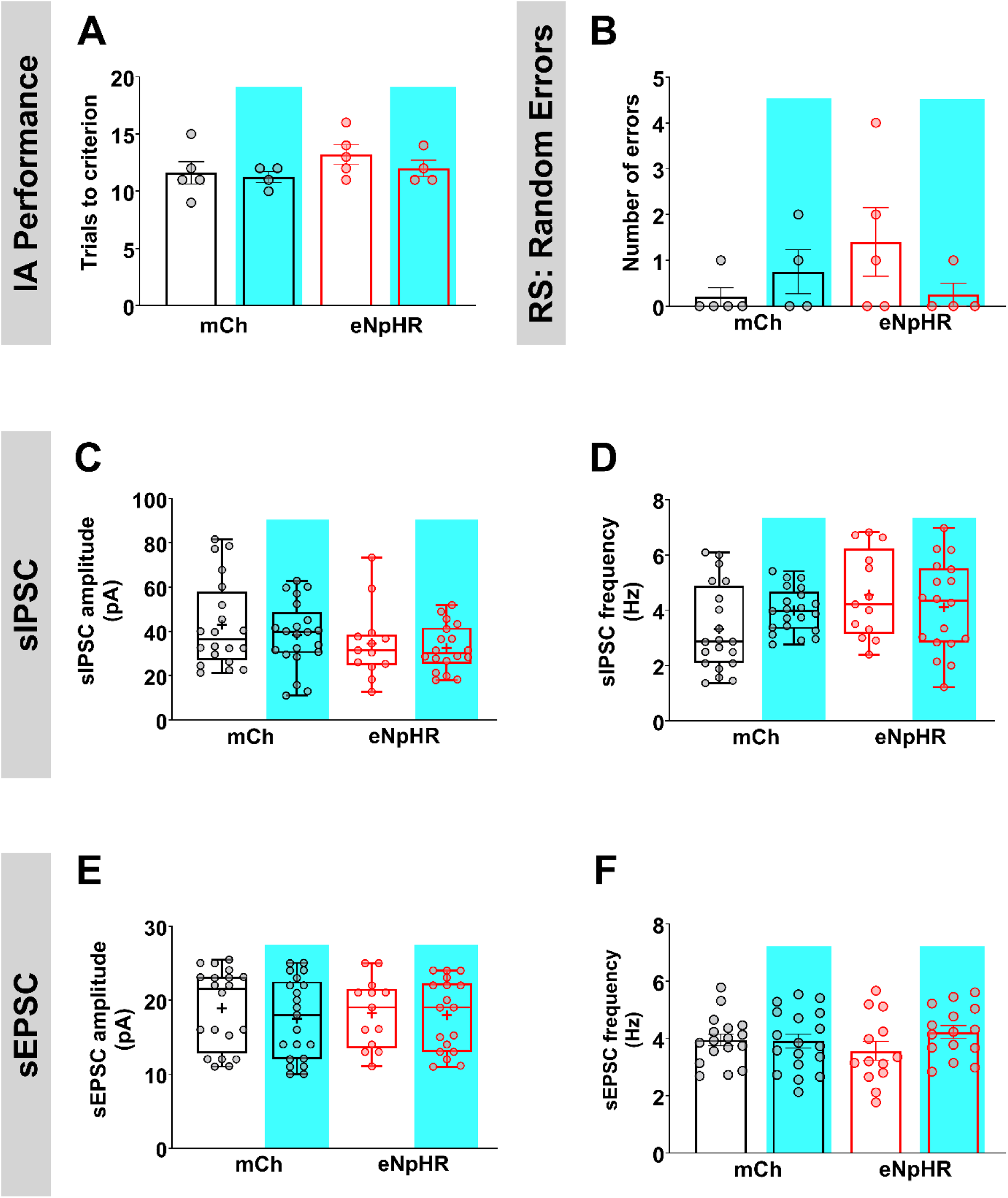
Additional behavioral measures and spontaneous synaptic recordings related to Figure 3. **(A-B)** There was no difference in learning of the IA or number of random (i.e., non-perseverative) errors during the rule-shift on Day 2. **(C–F)** sIPSC and sEPSC were recorded for 30–90 seconds from a subset of connected PFC-MD neurons. Sample sizes: mCh, no stim: N = 20 cells, n = 5 mice; mCh, 40 Hz: N = 21 cells, n = 4 mice; eNpHR, no stim: N = 13 cells, n = 5 mice; eNpHR, 40 Hz: N = 18 cells, n = 4 mice. **(C)** sIPSC amplitudes did not significantly differ among groups (Kruskal-Wallis test, p = 0.24). **(D)** sIPSC frequency also showed no significant differences (Kruskal-Wallis test, p = 0.14). **(E–F)** Similar analyses were performed for sEPSCs. **(E)** sEPSC amplitude was unchanged among groups (Kruskal-Wallis test, p = 0.74). **(F)** sEPSC frequency also showed no significant effects (main effect of eNpHR: p = 0.95; 40 Hz: p = 0.35; interaction: p = 0.28).

**Fig. S4.**
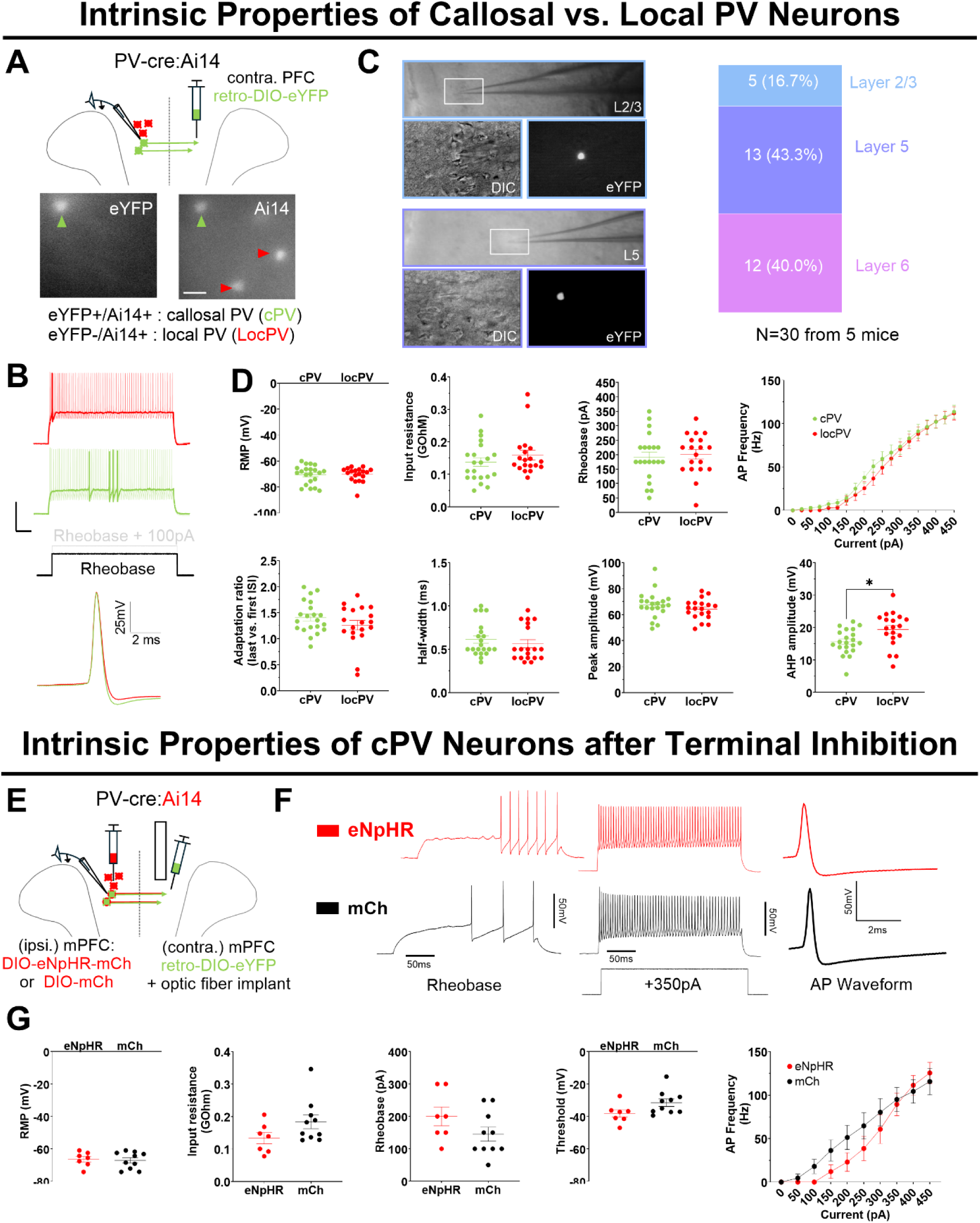
Intrinsic properties of callosally– and locally-projecting PV neurons and the effect of callosal PV terminal inhibition. **(A, top)** Experimental design. Callosally-projecting (cPV) neurons were retrogradely labeled in one mPFC by injecting AAVrg-DIO-eYFP into the contralateral mPFC of *PV-Cre:Ai14* mice. Whole-cell current-clamp recordings were performed to compare intrinsic properties between cPV and locally-projecting PV (locPV) neurons which were eYFP+/tdTomato+ or eYFP-/tdTomato+, respectively. **(A, bottom)** Representative images showing locPV (eYFP-/tdTomato+) and cPV (eYFP+/tdTomato+) neurons. Scale bar: 12.5 μm. **(B, top)** Example current-clamp responses of cPV neurons (green) and locPV neurons (red) to depolarizing current injections at rheobase (bold) and rheobase +100 pA. **(B, bottom)** Example action potential waveforms of cPV and locPV neurons, which were nearly indistinguishable. **(C, left)** Representative images of recorded cPV neurons in layer 2/3 and layer 5 of the medial prefrontal cortex (mPFC). Scale bars: 50 μm and 12.5 μm. **(C, right)** Laminar distribution of recorded cPV neurons (*N* = 30 from 5 mice). **(D)** cPV neurons exhibited overall similar intrinsic properties compared to locPV neurons. However, the afterhyperpolarization amplitude was significantly smaller in cPV neurons (15.7 ± 0.85 mV, *N* = 21, *n* = 5) compared to locPV neurons (19.4 ± 1.2 mV, *N* = 19, *n* = 5; unpaired *t* test, *p* < 0.05). **(E)** Experimental design. In *PV-Cre:Ai14* mice, we injected AAV5-DIO-eNpHR-mCherry (‘eNpHR,’ experimental group) or AAV5-DIO-mCherry (‘mCh,’ control group) into one mPFC, and AAVrg-DIO-eYFP into the contralateral mPFC. A 0.2 mm fiber optic was implanted targeting the PL/IL border contralateral to the side of virus injection for eNpHR / mCh. Mice performed the rule-shift (RS) task and the light for cPV terminal inhibition was delivered during the RS portion (as in Fig. 2–3), followed by current-clamp recordings from cPV neurons within 24 hours. **(F)** Example traces showing current-clamp responses at rheobase (left) and at a 350 pA current step (middle), and corresponding action potential waveforms (right) from cPV neurons after cPV terminal inhibition. **(G)** Inhibiting cPV axons during RS did not significantly alter the intrinsic properties of cPV neurons. Data are presented as mean ± SEM. See table S4 for detailed comparisons of intrinsic properties.

**Table S1.**
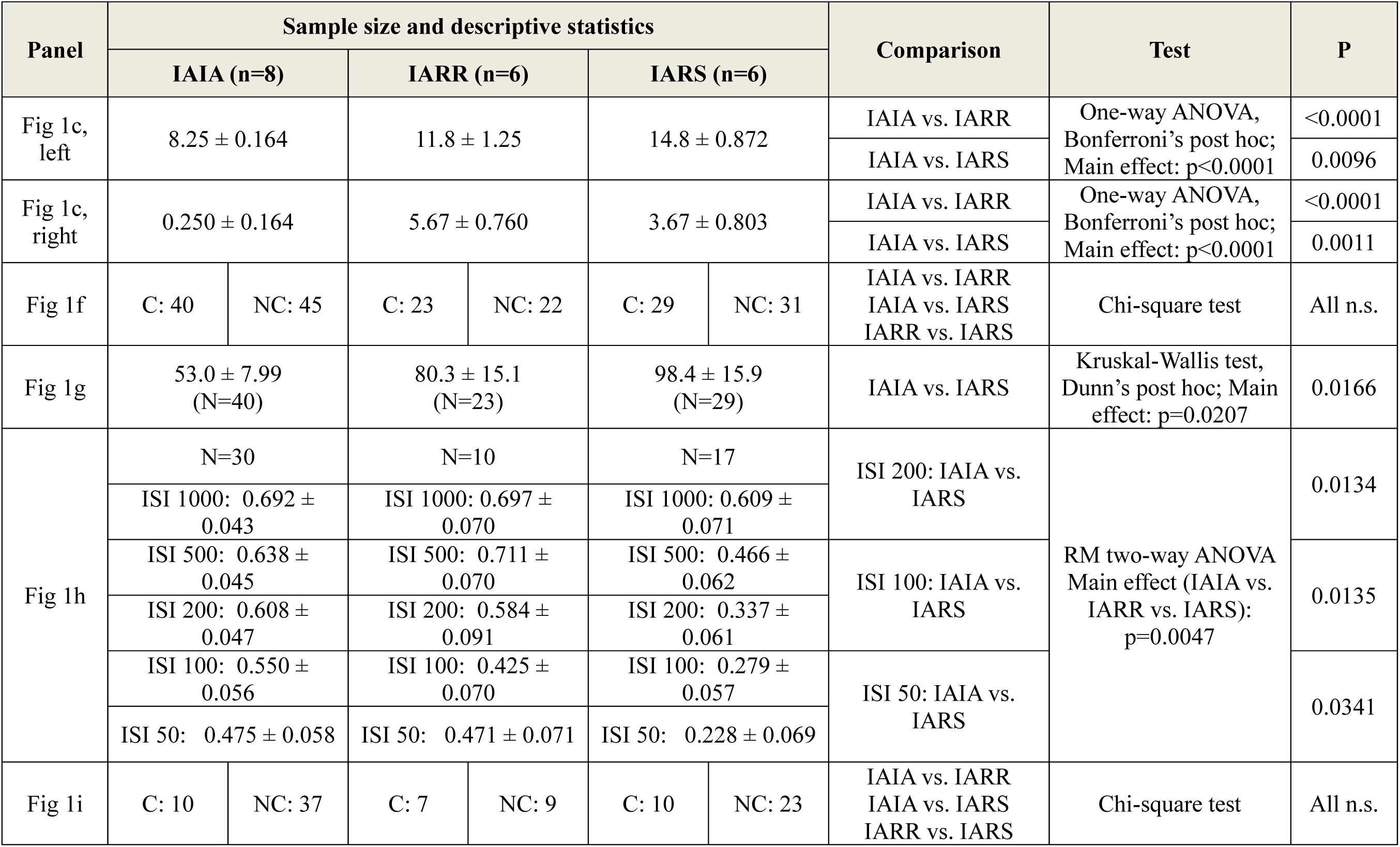
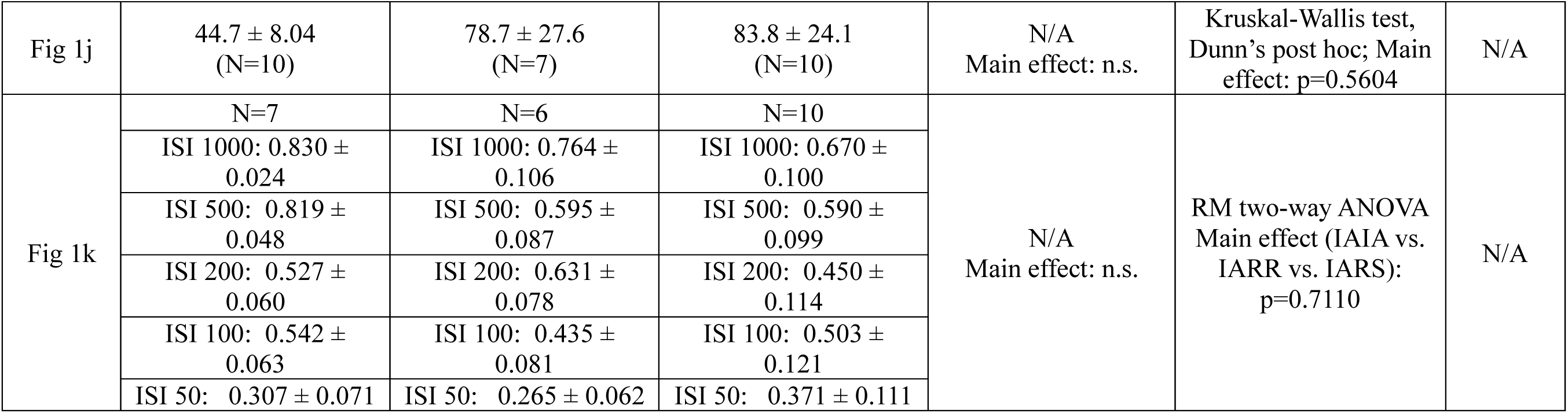
Summary of detailed statistics and additional oIPSC analysis related to Figure 1. C: connected; NC: not connected; ISI: inter-stimulus interval; n.s.: not significant.

| Panel | Sample size and descriptive statistics |  |  |  |  |  | Comparison | Test | P |
| --- | --- | --- | --- | --- | --- | --- | --- | --- | --- |
|  | IAIA (n=8) |  | IARR (n=6) |  | IARS (n=6) |  |  |  |  |
| Fig 1c, left | 8.25 ± 0.164 |  | 11.8 ± 1.25 |  | 14.8 ± 0.872 |  | IAIA vs. IARR | One-way ANOVA, Bonferroni's post hoc; Main effect: p<0.0001 | <0.0001 |
|  |  |  |  |  |  |  | IAIA vs. IARS |  | 0.0096 |
| Fig 1c, right | 0.250 ± 0.164 |  | 5.67 ± 0.760 |  | 3.67 ± 0.803 |  | IAIA vs. IARR | One-way ANOVA, Bonferroni's post hoc; Main effect: p<0.0001 | <0.0001 |
|  |  |  |  |  |  |  | IAIA vs. IARS |  | 0.0011 |
| Fig 1f | C: 40 | NC: 45 | C: 23 | NC: 22 | C: 29 | NC: 31 | IAIA vs. IARR<br>IAIA vs. IARS<br>IARR vs. IARS | Chi-square test | All n.s. |
| Fig 1g | 53.0 ± 7.99<br>(N=40) |  | 80.3 ± 15.1<br>(N=23) |  | 98.4 ± 15.9<br>(N=29) |  | IAIA vs. IARS | Kruskal-Wallis test, Dunn's post hoc; Main effect: p=0.0207 | 0.0166 |
| Fig 1h | N=30 |  | N=10 |  | N=17 |  | ISI 200: IAIA vs. IARS | RM two-way ANOVA<br>Main effect (IAIA vs. IARR vs. IARS):<br>p=0.0047 | 0.0134 |
|  | ISI 1000: 0.692 ± 0.043 |  | ISI 1000: 0.697 ± 0.070 |  | ISI 1000: 0.609 ± 0.071 |  |  |  |  |
|  | ISI 500: 0.638 ± 0.045 |  | ISI 500: 0.711 ± 0.070 |  | ISI 500: 0.466 ± 0.062 |  | ISI 100: IAIA vs. IARS |  | 0.0135 |
|  | ISI 200: 0.608 ± 0.047 |  | ISI 200: 0.584 ± 0.091 |  | ISI 200: 0.337 ± 0.061 |  |  |  |  |
|  | ISI 100: 0.550 ± 0.056 |  | ISI 100: 0.425 ± 0.070 |  | ISI 100: 0.279 ± 0.057 |  | ISI 50: IAIA vs. IARS |  | 0.0341 |
|  | ISI 50: 0.475 ± 0.058 |  | ISI 50: 0.471 ± 0.071 |  | ISI 50: 0.228 ± 0.069 |  |  |  |  |
| Fig 1i | C: 10 | NC: 37 | C: 7 | NC: 9 | C: 10 | NC: 23 | IAIA vs. IARR<br>IAIA vs. IARS<br>IARR vs. IARS | Chi-square test | All n.s. |

|  |  |  |  |  |  |  |
| --- | --- | --- | --- | --- | --- | --- |
| Fig 1j | 44.7 ± 8.04<br>(N=10) | 78.7 ± 27.6<br>(N=7) | 83.8 ± 24.1<br>(N=10) | N/A<br>Main effect: n.s. | Kruskal-Wallis test,<br>Dunn's post hoc; Main<br>effect: p=0.5604 | N/A |
| Fig 1k | N=7 | N=6 | N=10 | N/A<br>Main effect: n.s. | RM two-way ANOVA<br>Main effect (IAIA vs.<br>IARR vs. IARS):<br>p=0.7110 | N/A |
|  | ISI 1000: 0.830 ±<br>0.024 | ISI 1000: 0.764 ±<br>0.106 | ISI 1000: 0.670 ±<br>0.100 |  |  |  |
|  | ISI 500: 0.819 ±<br>0.048 | ISI 500: 0.595 ±<br>0.087 | ISI 500: 0.590 ±<br>0.099 |  |  |  |
|  | ISI 200: 0.527 ±<br>0.060 | ISI 200: 0.631 ±<br>0.078 | ISI 200: 0.450 ±<br>0.114 |  |  |  |
|  | ISI 100: 0.542 ±<br>0.063 | ISI 100: 0.435 ±<br>0.081 | ISI 100: 0.503 ±<br>0.121 |  |  |  |
|  | ISI 50: 0.307 ± 0.071 | ISI 50: 0.265 ± 0.062 | ISI 50: 0.371 ± 0.111 |  |  |  |

**Table S2.**
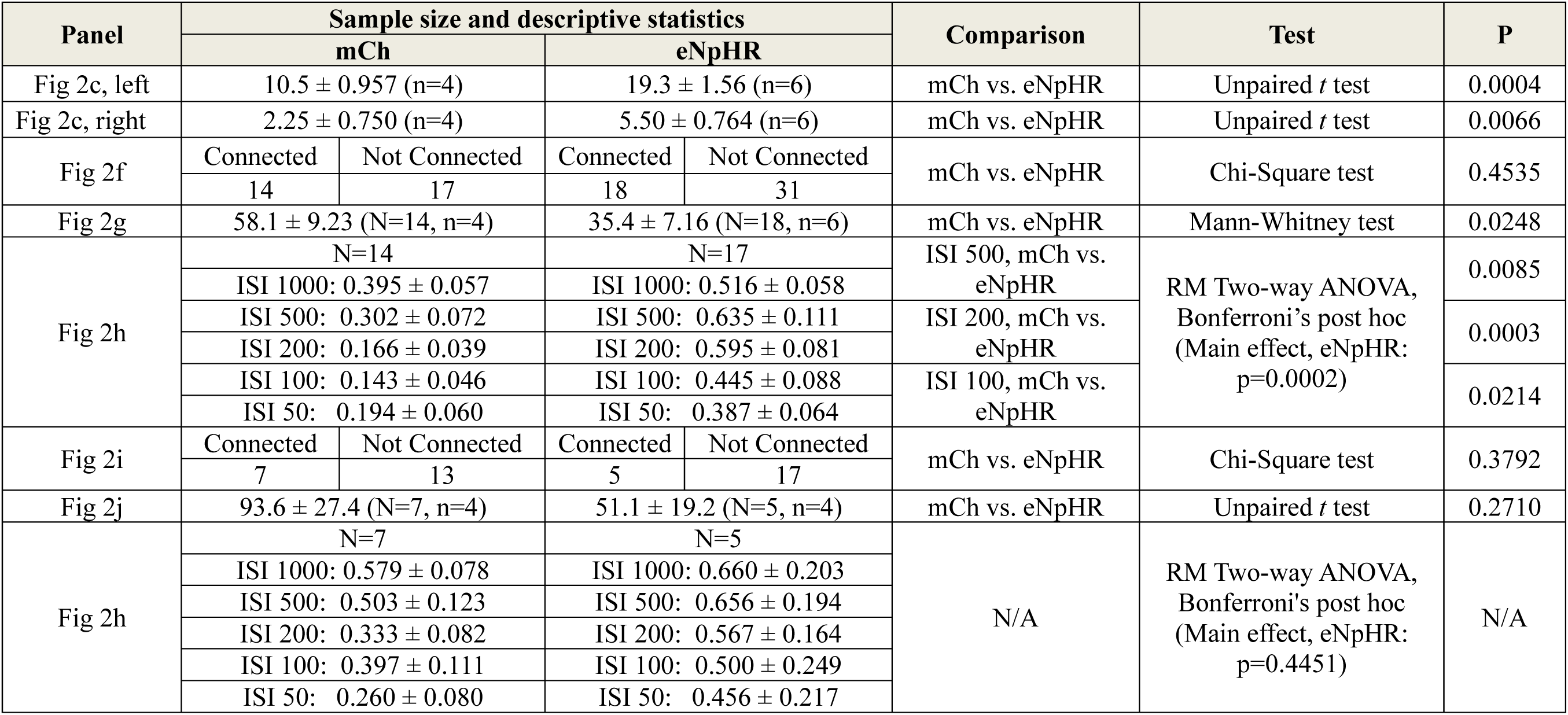
Summary of sample size and statistical comparisons in Figure 2.

**Table S3.** Summary of sample size and statistical comparisons in Figure 3.

| Panel | Sample size and descriptive statistics |  |  |  | Comparison | Test | P |
| --- | --- | --- | --- | --- | --- | --- | --- |
|  | mCh |  | eNpHR |  |  |  |  |
| Fig 3c, left | 11.9 ± 0.633 (n=9) |  | 21.9 ± 2.16 (n=9) |  | mCh vs. eNpHR | Unpaired *t* test | 0.0004 |
| Fig 3c, right | 2.67 ± 0.408 (n=9) |  | 8.22 ± 1.49 (n=9) |  | mCh vs. eNpHR | Unpaired *t* test | 0.0066 |
| Fig 3d, left | D2, no stim (n=5)<br>11.8 ± 0.663 | D2, 40Hz (n=4)<br>10.8 ± 0.629 | D2, no stim (n=5)<br>18.0 ± 2.00 | D2, 40Hz (n=4)<br>11.0 ± 0.408 | D2, no stim, mCh vs. eNpHR | Two-way ANOVA, Bonferroni's post hoc<br>Main effect of eNpHR: p=0.0209<br>Main effect of D2, 40Hz: p=0.0058<br>Interaction: p=0.0308 | 0.0129 |
|  |  |  |  |  | eNpHR, D2, no stim vs. D2 40Hz |  | 0.0080 |
|  |  |  |  |  | eNpHR, D2, no stim vs. mCh, D2 40Hz |  | 0.0060 |
| Fig 3d, right | D2, no stim (n=5)<br>11.8 ± 0.663 | D2, 40Hz (n=4)<br>10.8 ± 0.629 | D2, no stim (n=5)<br>18.0 ± 2.00 | D2, 40Hz (n=4)<br>11.0 ± 0.408 | D2, no stim, mCh vs. eNpHR | Two-way ANOVA, Bonferroni's post hoc<br>Main effect, eNpHR: p=0.0039<br>Main effect, D2, 40Hz: p=0.0193<br>Interaction: p=0.0285 | 0.0035 |
|  |  |  |  |  | eNpHR, D2, no stim vs. D2, 40Hz |  | 0.0175 |
|  |  |  |  |  | eNpHR, D2 no stim vs. mCh, D2, 40Hz |  | 0.0044 |
| Fig 3e | D2, no stim (n=5)<br>Connected: N=20<br>Not Connected: N=12 | D2, 40Hz (n=4)<br>Connected: N=22<br>Not Connected: N=12 | D2, no stim (n=5)<br>Connected: N=13<br>Not Connected: N=30 | D2, 40Hz (n=4)<br>Connected: N=18<br>Not Connected: N=11 | D2, no stim, mCh vs. eNpHR | Chi-Square test | 0.0054 |
|  |  |  |  |  | eNpHR, D2, no stim vs. D2 40Hz |  | 0.0075 |
|  |  |  |  |  | eNpHR, D2, no stim vs. mCh, D2, 40Hz |  | 0.0026 |
| Fig 3f | D2, no stim (N=20, n=5)<br>103 ± 15.5 | D2, 40 Hz (N=22, n=4)<br>114 ± 14.1 | D2, no stim (N=13, n=5)<br>37.6 ± 5.39 | D2, 40 Hz (N=18, n=4)<br>102 ± 21.8 | D2, no stim, mCh vs. eNpHR | Kruskal-Wallis test, Dunn's post hoc<br>p=0.0020 | 0.0112 |
|  |  |  |  |  | eNpHR, D2, no stim vs. D2, 40Hz |  | 0.0478 |
|  |  |  |  |  | eNpHR, D2 no stim vs. mCh, D2 40Hz |  | 0.0013 |
| Fig 3g | D2, no stim<br>(N=19, n=5) | D2, 40 Hz<br>(N=19, n=4) | D2, no stim<br>(N=10, n=5) | D2, 40 Hz<br>(N=17, n=4) | N/A<br>Main effects: n.s | RM Two-way ANOVA<br>Main effect of eNpHR:<br>p=0.1798<br>Interaction: 0.7829 | N/A. |
|  | ISI 1000:<br>0.748 ± 0.073 | ISI 1000:<br>0.782 ± 0.072 | ISI 1000:<br>0.698 ± 0.130 | ISI 1000:<br>0.604 ± 0.065 |  |  |  |
|  | ISI 500:<br>0.673 ± 0.056 | ISI 500:<br>0.581 ± 0.071 | ISI 500:<br>0.524 ± 0.090 | ISI 500:<br>0.500 ± 0.074 |  |  |  |
|  | ISI 200:<br>0.609 ± 0.074 | ISI 200:<br>0.486 ± 0.060 | ISI 200:<br>0.468 ± 0.093 | ISI 200:<br>0.397 ± 0.063 |  |  |  |
|  | ISI 100:<br>0.519 ± 0.058 | ISI 100:<br>0.384 ± 0.053 | ISI 100:<br>0.356 ± 0.076 | ISI 100:<br>0.317 ± 0.065 |  |  |  |
|  | ISI 50:<br>0.354 ± 0.060 | ISI 50:<br>0.300 ± 0.054 | ISI 50:<br>0.244 ± 0.062 | ISI 50:<br>0.247 ± 0.054 |  |  |  |
| Fig 3h | D2, no stim<br>(n=5) | D2 40Hz (n=4) | D2 no stim<br>(n=5) | D2, 40Hz (n=4) | D2 no stim, mCh vs.<br>eNpHR | Chi-Square test | 0.4853 |
|  | Connected:<br>N=15 | Connected:<br>N=13 | Connected: N=8 | Connected:<br>N=6 | eNpHR, D2 no stim vs.<br>D2 40Hz |  | 0.2575 |
|  | Not Connected:<br>N=14 | Not Connected:<br>N=14 | Not Connected:<br>N=10 | Not Connected:<br>N=16 | eNpHR, D2 no stim vs.<br>mCh, D2 40Hz |  | 0.8073 |
| Fig 3i | D2, no stim<br>(N=15, n=5)<br>80.5 ± 15.9 | D2, 40 Hz<br>(N=13, n=4)<br>116 ± 16.8 | D2, no stim<br>(N=8, n=5)<br>37.7 ± 4.35 | D2, 40 Hz<br>(N=6, n=4)<br>71.1 ± 15.4 | D2 no stim, mCh vs.<br>eNpHR | Kruskal-Wallis test,<br>Dunn's post hoc<br>p=0.0211 | 0.1996 |
|  |  |  |  |  | eNpHR, D2 no stim vs.<br>D2 40Hz |  | 0.5370 |
|  |  |  |  |  | eNpHR, D2 no stim vs.<br>mCh, D2 40Hz |  | 0.0112 |
| Fig 3j | D2, no stim<br>(N=12, n=5) | D2, 40 Hz<br>(N=11, n=4) | D2, no stim<br>(N=6, n=5) | D2, 40 Hz<br>(N=5, n=4) | N/A | RM Two-way ANOVA<br>Main effect of eNpHR:<br>p=0.1344<br>Interaction: p=0.0749 | N/A |
|  | ISI 1000:<br>0.773 ± 0.081 | ISI 1000: 0.933<br>± 0.110 | ISI 1000: 0.526<br>± 0.121 | ISI 1000: 0.596<br>± 0.200 |  |  |  |
|  | ISI 500:<br>0.587 ± 0.069 | ISI 500: 0.741<br>± 0.061 | ISI 500: 0.384<br>± 0.161 | ISI 500:<br>0.455 ± 0.191 |  |  |  |
|  | ISI 200:<br>0.509 ± 0.050 | ISI 200: 0.716<br>± 0.050 | ISI 200: 0.391<br>± 0.146 | ISI 200: 0.453<br>± 0.188 |  |  |  |
|  | ISI 100:<br>0.515 ± 0.057 | ISI 100: 0.617<br>± 0.065 | ISI 100: 0.291<br>± 0.110 | ISI 100: 0.448<br>± 0.185 |  |  |  |
| | ISI 50:<br>$0.367 \pm 0.074$ | ISI 50: $0.411 \pm 0.062$ | ISI 50: $0.450 \pm 0.221$ | ISI 50: $0.556 \pm 0.171$ | | | |

**Table S4.** Summary of intrinsic electrophysiological properties of mPFC LocPV and cPV neurons (related to Fig. S4). RMP: resting membrane potential; ISI: inter-spike interval; AHP: after-hyperpolarization.

| (Related to Fig. S4D) Intrinsic properties of cPV vs. locPV neurons |  |  |  |  |
| --- | --- | --- | --- | --- |
| Measure | Descriptive Statistics |  | Test | P |
|  | cPV (N=21, n=5) | locPV (N=19, n=5) |  |  |
| RMP (mV) | $-70.5 \pm 1.55$ | $-70.2 \pm 1.25$ | Unpaired *t* test | 0.8932 |
| R<sub>input</sub> (GOhm) | $0.138 \pm 0.0134$ | $0.159 \pm 0.0150$ | Unpaired *t* test | 0.2903 |
| Time constant (ms) | $6.32 \pm 0.880$ | $10.6 \pm 2.43$ | Mann-Whitney test | 0.1607 |
| Rheobase (pA) | $192 \pm 17.5$ | $201 \pm 16.4$ | Unpaired *t* test | 0.6917 |
| AP threshold (mV) | $-28.0 \pm 3.29$ | $-35.3 \pm 2.54$ | Unpaired *t* test | 0.0904 |
| F/I relationship | N/A | N/A | 2-way RM ANOVA;<br>Factor: cPV vs. locPV | 0.4949 |
| Adaptation ratio<br>(last to first ISI) | $1.41 \pm 0.0672$ | $1.26 \pm 0.0895$ | Unpaired *t* test | 0.1888 |
| AP halfwidth (ms) | $0.612 \pm 0.0435$ | $0.563 \pm 0.0466$ | Unpaired *t* test | 0.4489 |
| AP peak amplitude<br>(mV) | $67.4 \pm 2.21$ | $64.1 \pm 1.81$ | Unpaired *t* test | 0.2667 |
| AHP amplitude<br>(mV) | $15.7 \pm 0.845$ | $19.4 \pm 1.23$ | Unpaired *t* test | 0.0133 |
| (Related to Fig. S4G) Intrinsic properties of cPV neurons after cPV terminal inhibition during RS |  |  |  |  |
| Measure | Descriptive statistics |  | Test | P |
|  | eNpHR (N=7, n=2) | mCh (N=10, n=2) |  |  |
| RMP (mV) | $-66.4 \pm 1.70$ | $-67.1 \pm 1.65$ | Unpaired *t* test | 0.7737 |
| R<sub>input</sub> (GOhm) | $0.133 \pm 0.0173$ | $0.184 \pm 0.0212$ | Unpaired *t* test | 0.1036 |
| Time constant (ms) | $12.3 \pm 3.55$ | $10.8 \pm 2.12$ | Mann-Whitney test | 0.6691 |
| Rheobase (pA) | $200 \pm 28.9$ | $145 \pm 21.7$ | Unpaired *t* test | 0.1407 |
| AP threshold (mV) | $-38.1 \pm 2.28$ | $-31.5 \pm 2.29$ | Unpaired $t$ test | 0.4584 |
| F/I relationship | N/A | N/A | 2-way RM ANOVA;<br>Factor: mCh vs. eNpHR | 0.4281 |
| Adaptation ratio<br>(last to first ISI) | $1.340 \pm 0.147$ | $1.316 \pm 0.125$ | Unpaired $t$ test | 0.9022 |
| AP halfwidth (ms) | $0.529 \pm 0.0721$ | $0.585 \pm 0.0522$ | Unpaired $t$ test | 0.5441 |
| AP peak amplitude<br>(mV) | $61.94 \pm 3.88$ | $65.80 \pm 2.74$ | Unpaired $t$ test | 0.4315 |
| AHP amplitude<br>(mV) | $18.09 \pm 1.77$ | $18.46 \pm 1.25$ | Unpaired $t$ test | 0.8735 |

## Notes

### Competing Interest Statement

The authors have declared no competing interest.

